# Curvelet Transform-based Sparsity Promoting Algorithm for Fast Ultrasound Localization Microscopy

**DOI:** 10.1101/2022.02.02.478910

**Authors:** Qi You, Joshua D. Trzasko, Matthew R. Lowerison, Xi Chen, Zhijie Dong, Nathiya Vaithiyalingam ChandraSekaran, Daniel A. Llano, Shigao Chen, Pengfei Song

## Abstract

Ultrasound localization microscopy (ULM) based on microbubble (MB) localization was recently introduced to overcome the resolution limit of conventional ultrasound. However, ULM is currently challenged by the requirement for long data acquisition times to accumulate adequate MB events to fully reconstruct vasculature. In this study, we present a curvelet transform-based sparsity promoting (CTSP) algorithm that improves ULM imaging speed by recovering missing MB localization signal from data with very short acquisition times. CTSP was first validated in a simulated microvessel model, followed by the chicken embryo chorioallantoic membrane (CAM), and finally, in the mouse brain. In the simulated microvessel study, CTSP robustly recovered the vessel model to achieve an 86.94% vessel filling percentage from a corrupted image with only 4.78% of the true vessel pixels. In the chicken embryo CAM study, CTSP effectively recovered the missing MB signal within the vasculature, leading to marked improvement in ULM imaging quality with a very short data acquisition. Taking the optical image as reference, the vessel filling percentage increased from 2.7% to 42.2% using 50ms of data acquisition after applying CTSP. CTSP used 80% less time to achieve the same 90% maximum saturation level as compared with conventional MB localization. We also applied CTSP on the microvessel flow speed maps and found that CTSP was able to use only 1.6s of microbubble data to recover flow speed images that have similar qualities as those constructed using 33.6s of data. In the mouse brain study, CTSP was able to reconstruct the majority of the cerebral vasculature using 1-2s of data acquisition. Additionally, CTSP only needed 3.2s of microbubble data to generate flow velocity maps that are comparable to those using 129.6s of data. These results suggest that CTSP can facilitate fast and robust ULM imaging especially under the circumstances of inadequate microbubble localizations.

## I. INTRODUCTION

Super-resolution ultrasound microvessel imaging, based on microbubble (MB) localization, is a rapidly growing field. Inspired by the optical localization microscopy super-resolution techniques such as fPALM and STORM [1–2], Desailly et al. [3], Viessmann et al. [4], and O’Reilly et al. [5] conducted several early studies on super-resolution ultrasound imaging using injected MBs as acoustically localizable point sources to overcome the acoustic diffraction limit. More recent works from Christensen-Jeffries et al. [6] and Errico et al. [7] demonstrated successful applications using MB localization and tracking techniques, collectively named ultrasound localization microscopy (ULM), to image *in-vivo* microvasculature structures as well as map blood velocity at a super-resolution scale. Various follow up studies have quickly emerged to improve this technique, including MB signal denoising to reduce the localization of noise and improve the accuracy of localization [8], motion correction to account for sub-wavelength tissue movement [9], improvements in localization techniques and accuracy [10], and using high-order statistic model to enhance spatial resolution [11].

Despite the great potential of ULM in a wide range of preclinical and clinical applications, its practical value is still challenged by several notable limitations such as the requirement for long-duration data acquisitions [12–13] and the expensive and complex post-processing algorithms involving MB localization and tracking [14–15]. To achieve successful ULM with accurate vascular reconstruction, it is necessary to accumulate ample amount of spatially isolated MB events to facilitate robust localization. This requirement inevitably imposes a trade-off between imaging speed and localization accuracy: lowering the MB concentration by dilution results in less MB signal overlap and higher localization accuracy, but not without the cost of a longer required data acquisition time to ensure complete MB perfusion of the microvasculature; by comparison, higher MB concentrations lead to faster MB saturation and imaging speed, but the increased amount of superimposed MB signal are challenging for localization. Pragmatically, since physiological movements (e.g. breathing) and operator-induced motions (e.g., free-hand scanning in the clinic) are common and inevitable, a faster imaging speed with shorter data acquisition time is essential for successful *in-vivo* implementation of ULM. As such, developing an image reconstruction algorithm that can recover the microvasculature by utilizing a short data acquisition with a small amount of MB events is critical for widespread use of ULM.

To this end, our groups have introduced several techniques that include MB separation post-processing [16] and Kalman filter-based inpainting [17] to reduce ULM data acquisition times and improve ULM imaging speed. The MB separation technique [16] designed a set of 3D Fourier domain filters to separate MB events with different spatiotemporal characteristics (e.g., movement speed, flow direction) to improve MB localization and tracking performance at high MB concentrations. However, this method aggravates the computational burden of ULM due to the requirement of independent localization and tracking of MB events for each filter bandwidth. The Kalman filter-based inpainting method [17] generates smooth and continuous microvascular images by imposing constraints to MB movement and utilizing interpolation techniques to recover missing vessel signal. However, this approach may be limited to deal with complex flow patterns and tortuous vessels where MB movement trajectory becomes abrupt and unpredictable. In addition, a group of deep-learning-based ULM methods [18–22] have been developed recently to improve the localization accuracy and reconstruction speed. However, the deep-learning-based method works as a black box and its performance is highly related to the training dataset, which is usually difficult to collect from in-vivo applications. Besides, a recent superresolution technique SUSHI [23] was developed to improve the resolution of contrast enhanced ultrasound imaging by exploiting the sparsity of the microvessel signal in spatial domain. Inspired by these preceding studies, we discovered an algorithm to improve the ULM imaging speed by using only a small number of MBs.

For ULM, the reconstruction of a microvasculature from an inadequate amount of MB locations is related to the topic of “reconstruction from highly incomplete information” or “compressive sampling” in the field of image and signal processing [24–30]. The principle of compressive sampling has been successfully applied in the field of seismic and medical imaging, including magnetic resonance imaging (MRI) and X-ray tomography with significantly under-sampled source signal [26-27,30]. Similarly, in ULM, the isolated MB locations can be considered as spatial samples of a continuous microvessel structure. Sparse MB events caused by low MB concentrations or short acquisition times result in under-sampling of the vasculature, which can be recovered using the same principles as in compressive sampling theory.

To implement the compressive sampling theory in ULM, three components need to be pre-defined: a sparsifying transform, a random down-sampling strategy that confines aliasing, and an effective recovery algorithm that promotes sparsity in the transform domain. The existence of MB events in the microvessels can be assumed to follow a Poisson distribution [18–19] and thus satisfies the random downsampling requirement. Many studies [25-26,29-34] have proved that recovery from compressive sampling can be achieved by minimizing the *ℓ*_1_ norm of the transformed signal, subjected to an *ℓ*_2_ norm data fidelity term. For the sparsifying transform, many signal reconstruction studies [26,29,33-37] based on compressive sampling chose multi-resolution analysis (MRA) methods such as wavelet or curvelet transforms. Among these MRA methods, the curvelet transform [37–42] attains high compression on curve-shaped structures due to the large correlations between curvelets and curved fronts. In addition, the sparsity-based super-resolution technique SUSHI [23] proposed a sparsifying transform based on the assumption of spatially sparse microvessel signal. Inspired by these previous studies, we propose a microvessel reconstruction method that promotes the sparsity both in the curvelet transform domain and the spatial domain, aiming to recover the microvessel structure from under-sampled MB positions while maintaining the spatial features of the microvessels.

The rest of the paper is organized as follows. In Section II, we present the principle of curvelet transform-based sparsity promoting (CTSP) algorithm, followed by the experiment setup which includes the *in-silico* study, *in-vivo* chorioallantoic membrane (CAM) study, and *in-vivo* mouse brain study. In Section III, the results of all experiments are presented. In Section IV, we finalize the paper with discussion and conclusions.

## II. Materials and Methods

### A. Principle of CTSP

#### 1) Curvelet transform

The curvelet transform was first introduced in the field of multi-resolution analysis by Candès *et al* [37–42]. Similar as the wavelet and Fourier transforms, the curvelet transform decomposes a signal in spatial domain into a combination of a series of signal bases, namely curvelets. The curvelets, notated as *φ_j,l,k_*(*x*), are a series of waveforms defined with three parameters: scale *j*, orientation *l*, and translation *k*. To detail the mathematical definition of curvelets, we start by defining a pair of nonnegative and real-valued windows *W*(*r*), *r* ∈ (1/2,2) and *V*(*t*), *t* ∈ [-1,1]. A 2D curvelet *φ_j_*(*x,y*) with scale *j* is defined in the Fourier domain as

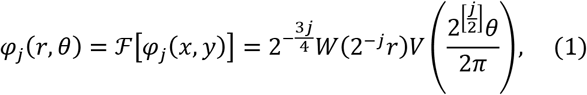

where (*r, θ*) represent the coordinates in the polar system, 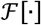 represents the 2D Fourier transform, [*j*/2] represents the integer part of *j*/2. Then we define a sequence of rotation angles as

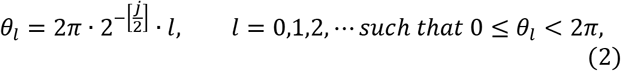

and a rotation matrix as

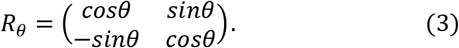

Finally, we define a sequence of position vectors as

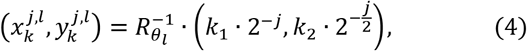

where *k* = (*k*_1_, *k*_2_) represents the translation vector in the 2D space. With the above notations, we can define the 2D curvelet with scale *j*, orientation *l* and translation *k* as:

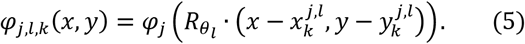

With the above definition of the 2D curvelet, a typical 2D curvelet transform can be defined with the following equation:

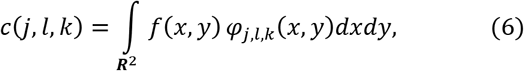

where *f*(*x, y*) represents the 2D signal in spatial domain with function *f* ∈ ***R***^2^. The definition of curvelets is further illustrated with a simple 64×64 2D space case as shown in Fig. 1. Take the curvelet *φ*_2,1_ in the red area in Fig. 1(a) as an example: its Fourier transform *φ*_2,1_(*r, θ*) is defined as the red area in Fig. 1(a) and its waveform is shown as a wedged wave in Fig. 1(e-g). When the scale parameter *j* increases, the curvelet will move to the outside layer in Fourier domain as shown in the green area in Fig. 1(a) and will become sharper in the spatial domain as shown in Fig. 1(b-d). When the orientation parameter *l* changes, the curvelet will rotate in both the Fourier domain and the spatial domain as shown in the blue area in Fig. 1(a) and Fig. 1(h-j). When the translation parameter *k* changes, the curvelet waveform will move to the corresponding position in 2D space. The curvelet coefficient *c*(*j, l, k*) is therefore the inner product or the correlation coefficient between the 2D spatial signal and the corresponding curvelet coefficient *φ_j,l,k_*(*x, y*).

**Figure 1.**
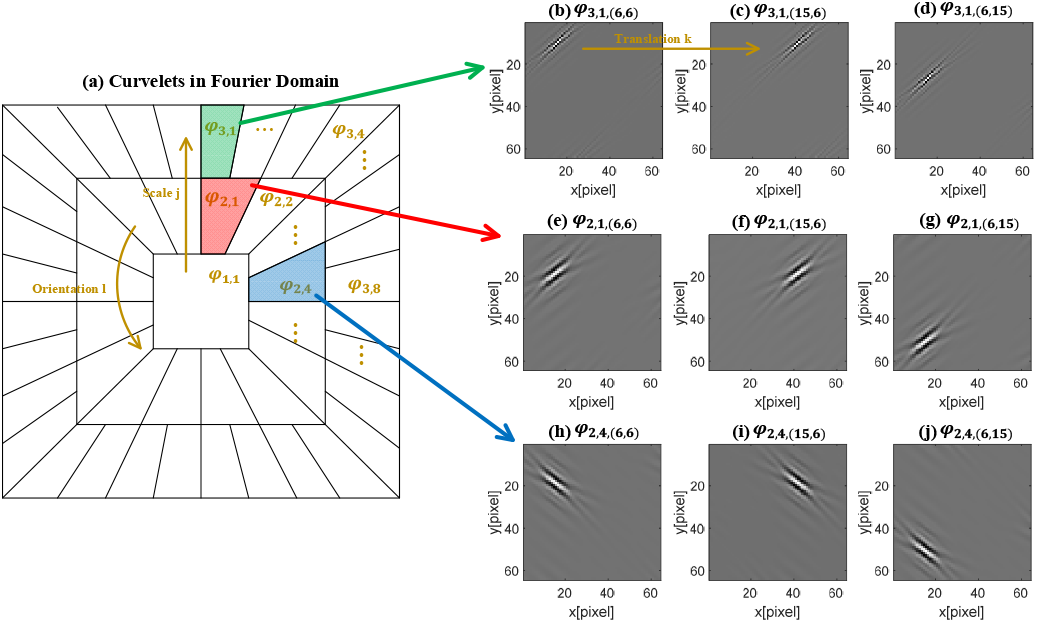
Curvelets defined in a 64×64 2D space. (a) Curvelets in Fourier domain with different supports. (b-j) Curvelets waveforms in spatial domain with different scale, orientation, and translation.

An important feature of the curvelet transform is the introduction of an orientation parameter into the curvelet bases. This is demonstrated with a simulated vessel example in Fig. 2 which has been reconstructed with a subset of curvelet coefficients. As shown in the magnified local region in Fig. 2(a), when the curvelet waveform has a similar orientation with the curved structure, the curvelet coefficient becomes larger due to the rising correlations whereas the curvelet coefficient will decrease rapidly when the orientation of the curvelet and the direction of the curved structure in the image do not match. Fig. 2(b-d) demonstrates that a few of the largest curvelet coefficients contain most of the structural information in the spatial domain, which guarantees the sparsity of the vesselshaped structure in the curvelet domain.

**Figure 2.**
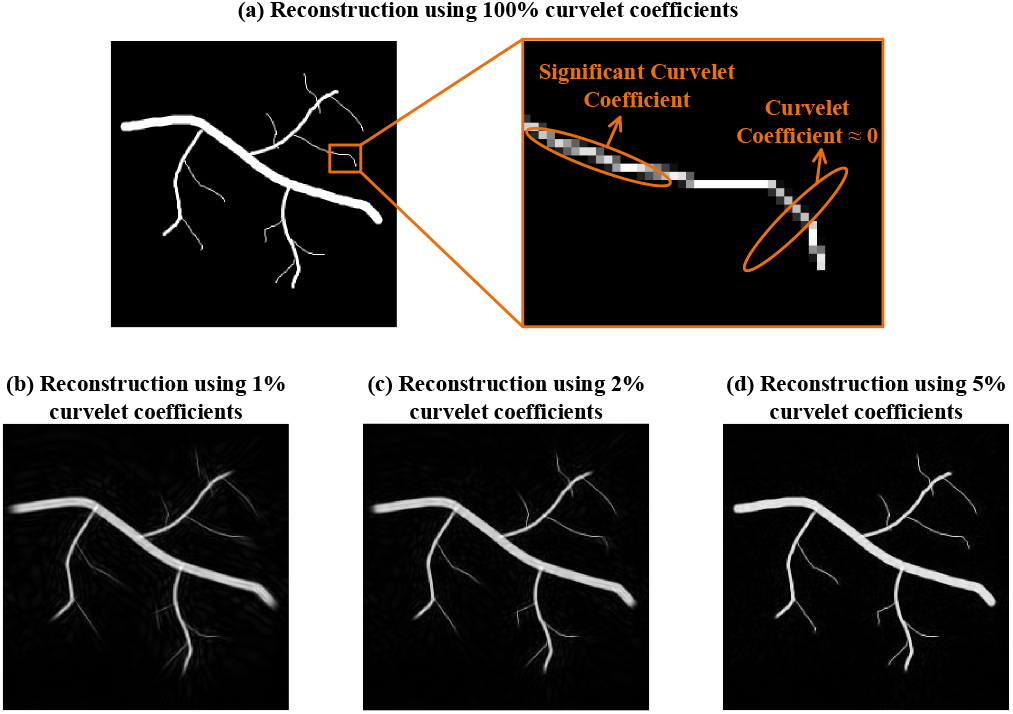
A simulated vessel reconstructed from (a) 100% of the largest curvelet coefficients (b) 1% of the largest curvelet coefficients (c) 2% of the largest curvelet coefficients (d) 5% of the curvelet coefficients. The orange ovals in (a) represent the shapes of two curvelet waveforms with different orientations.

#### 2) Compressive sampling-based ULM

The MB localization-based ULM can be described as a compressive sampling model:

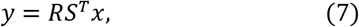

where *x* denotes the curvelet coefficients of the true vessel intensity map or velocity map, *S^T^* denotes the inverse curvelet transform, *R* denotes the spatial sampling matrix, and *y* denotes the corrupted vessel intensity map or velocity map. Eq. (7) represents the vessel intensity map generated from MB localization, or the velocity map generated from MB tracking trajectories, as the compressive spatial sampling of true intensity or velocity map. To recover the true vessels *x* from the corrupted vessel intensity or velocity map *y*, an *ℓ*_1_ norm minimization problem is formed to promote sparsity in the curvelet domain [25,31]:

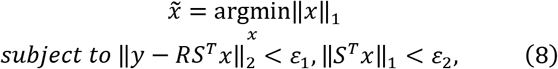

where *ε*_1_ represents the *ℓ*_2_ error in the measurement and *ε*_2_ represents the threshold to constrain the sparsity of the vessel structure in the spatial domain. The constrained optimization problem in Eq. (8) can be replaced by an unconstrained penalized optimization [26,29]:

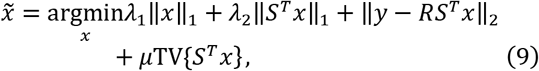

where *λ*_1_ and *λ*_2_ are regularization parameters associated with *ε*_1_ and *ε*_2_. TV{·} represents the total variation (TV) penalty term, which is used for suppressing ringing artifacts and recovering sharp edges [38]. Equation (8) can be iteratively solved by using the block-coordinate-relaxation (BCR) method. The details of the iterative algorithm are shown in Table I. Similar approaches have been used to solve the optimization problems with multiple data fidelity terms or penalty terms in the field of image inpainting [29], seismic wave recovery [26], and MRI reconstruction [44].

**TABLE I.**
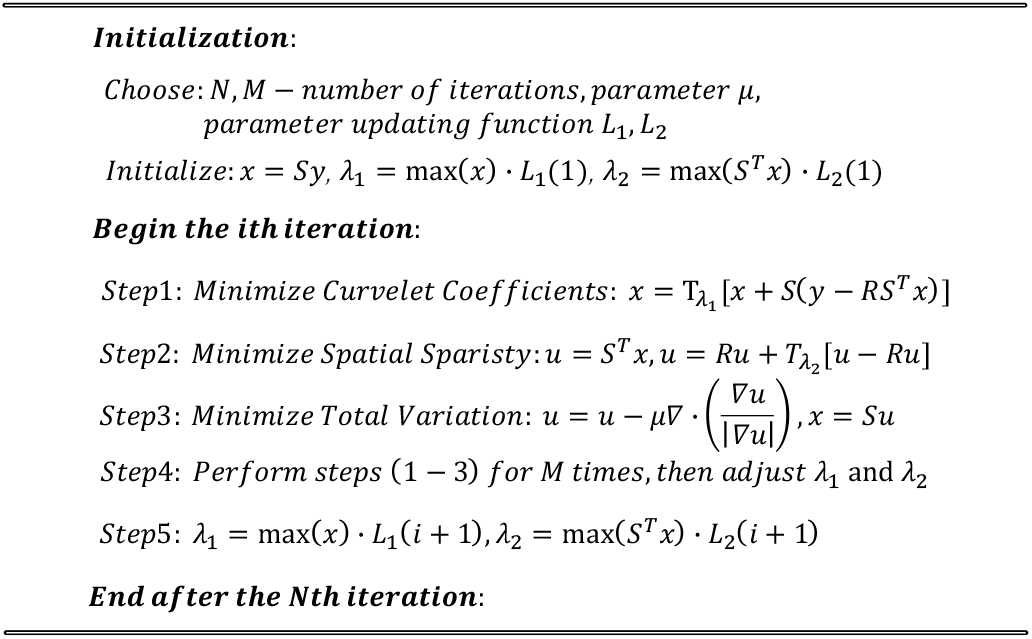
BLOCK-COORDINATE-RELAXATION WITH ITERATIVE THRESHOLD.

As shown in Table I, the BCR method splits Eq. (9) into three separate parts and solves them independently using three steps. Step 1 updates *x* with:

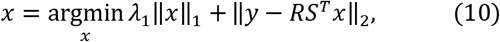

and is solved by using the soft thresholding [25]:

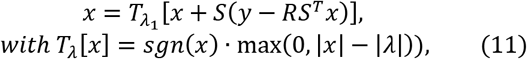

where *T_λ_* represents the soft threshold operator and *sgn*(*x*) represents the signum function. Step 2 first reconstructs the spatial image *u* from the curvelet coefficients *x* by *u* = *S^T^x* and then updates *u* with:

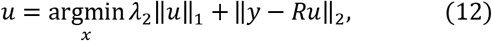

and can be also solved by the soft thresholding:

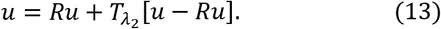

Step 3 minimizes the TV term by gradient descent [43] and then calculate the curvelet coefficients by implementing the curvelet transform *x* = *Su*. The regularization parameter *μ* is empirically chosen as a moderate fixed value that guarantees convergence. Step 4 performs steps (1-3) for M times, where M is the iteration number that controls the convergence of *x* under the current parameters *λ*_1_ and *λ*_2_. Then step 5 updates the parameters *λ*_1_ and *λ_2_* using the pre-defined updating functions *L*_1_ and *L*_2_. All 5 steps compose one loop of iteration and are performed N times, where the iteration number N is chosen to allow the algorithm to achieve a practically useful reconstruction.

In the proposed CTSP algorithm, the regularization parameters *λ*_1_ and *λ*_2_ control the soft thresholding for curvelet sparsity and spatial sparsity separately in every iteration. The parameter updating functions *L*_1_ and *L*_2_ represent the updating strategy of regularization parameters *λ*_1_ and *λ_2_*. In other studies [26,29], a common strategy is to design the updating function *L* as a monotonically decreasing function to shrink curvelet coefficients faster and generate more spatial connections in the beginning. As *L*_1_ lowers *λ*_1_, the convergence becomes slower and more accurate. A simple linear decreasing function used in the image inpainting study [29] is:

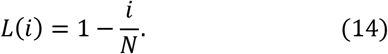

Where *i* represents the iteration index and *N* represents the number of iterations. In our study, we added several modifications to this linear model. First, the function *L*_2_ needs to be an increasing function as we want the spatial sparsity to be gradually promoted because the shrinkage of curvelet coefficients introduces an increasing amount of ringing artifacts during the iterations. Second, an adaptive amplitude parameter is required to adjust the increasing or decreasing rate of *L*_1_ or *L*_2_ based on different initial images. Based on these requirements, we designed the regularization parameter updating functions *L*_1_ and *L*_2_ as:

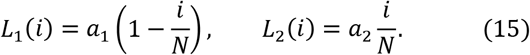

Here, the amplitude parameters *a*_1_ and *a*_2_ are chosen based on the initial images of specific applications. In our study, we first selected several ULM vessel images using data with different acquisition times and then empirically determined the parameters *a*_1_ and *a*_2_ to allow CTSP recovery to achieve the visually optimal reconstructed images. Then, a power regression is applied to fit the functions of *a*_1_ and *a*_2_ with the average pixel intensity of the initial image:

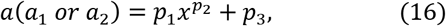

where *x* is the average pixel intensity of the initial ULM vessel image and (*p*_1_, *p*_2_, *p*_3_) are the power regression parameters. Together, equations (15-16) describe the procedure of determining the adaptive regularization parameter updating functions based on the initial ULM image to be recovered by CTSP.

The whole procedure of the CTSP algorithm is shown in Table I. There are five steps to implement in each iteration. Step 1 shrinks the curvelet coefficients using the current regularization parameter *λ*_1_. After step 1, the zero pixels around the isolated MB positions are filled, which recovers the missing vessel signal and generates ringing artifacts near the vessel boundaries. Step 2 minimizes the spatial sparsity using the regularization parameter *λ*_2_. This step resets some of the newly filled pixels based on *λ*_2_, which is proportional to the maximum absolute intensity of the current vessel map. Step 3 further adjusts the pixel intensities near the vessel boundaries to reduce the blurring by minimizing the total variation. After steps 2 and 3, the connection within the vessels is preserved while the artifacts around the vessel boundaries are removed. Step 4 repeats Steps 1-3 for M times to achieve a local convergence based on current *λ*_1_ and *λ*_2_. Finally Step 5 updates *λ*_1_ and *λ*_2_ based on the current recovered vessel map for the next iteration.

The performance of the proposed CTSP algorithm is subject to several factors: first, noise and false MB localizations can lead to compromised CTSP recovery. Applying a small median filter (e.g., 2 × 2) to the CTSP recovered image can mitigate this issue without blurring. Second, under the circumstances of high MB accumulations within dense microvessels (i.e. capillaries), CTSP performance may deteriorate because the assumption of sparsity for MB locations is violated, making the recovery an ill-posed problem. Therefore, for data with long durations of MB accumulation, it is more appropriate to apply CTSP to short time segments of MB data, and then accumulate all the CTSP recovered images on the short time segments to generate the final image.

### B. In-silico study

The performance of the proposed CTSP algorithm was first tested on a simple simulated vessel model. As shown in Fig. 3(a), the simulated vessel structure includes one artery and several branches with radii ranging from 2 to 8 pixels. The MB events at every pixel within the simulated vessel structure can be modelled as independent and identical Poisson distributions [18–19], where the Poisson rate parameter is proportional to the data acquisition time. The Poisson rate parameter does not affect the performance of CTSP. In our study, we assumed that the expected number of MB events per pixel and per unit time was 0.05. This number was selected based on the experimental observation, which nearly equals to the vessel filling percentage of conventional localization using 100ms acquisition in the CAM study. MB localization maps with 1 to 100 units of acquisition time were simulated, and then the true vessels were recovered using the proposed CTSP algorithm from each MB localization image. The curvelet transform in the CTSP algorithm is performed using the Curvelab toolbox provided by Candès et al [42].

**Figure 3.**
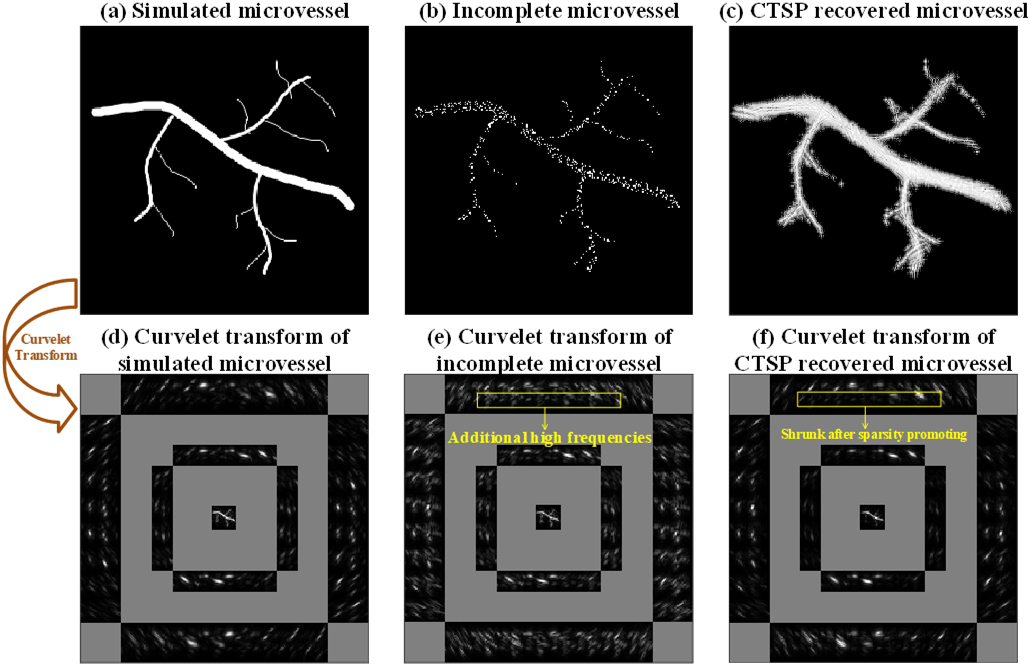
(a) Simulated microvessel image; (b) Corrupted image with random downsampling following a Poisson distribution (80.96% pixels removed); (c) CTSP recovered image; (d-f) Corresponding curvelet transforms of the images in (a-c).

To evaluate the performance of the proposed CTSP algorithm, we used the metric vessel filling percentage (*VF_percentage_*) and vessel filling precision (*VF_precision_*) defined as:

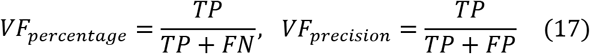

where true positives (TP) are pixels that are identified as true vessels in both the CTSP recovered image and the reference ground truth image, false negatives (FN) are pixels that are identified as non-vessels in the CTSP recovered image but are true vessel pixels in the reference ground truth image, and false positive (FP) are pixels that are identified as vessels in the CTSP recovered image but are non-vessels in the ground truth image. A 40 dB dynamic range threshold was used in the CTSP recovered image to distinguish vessel and non-vessel pixels. *VF_percentage_* measures the percentage of the vessels that are correctly reconstructed. *VF_precision_* measures the percentage of the recovered vessels that are true vessels.

### C. In-vitro flow phantom study

An *in-vitro* flow phantom model was used to validate the velocity recovery using CTSP algorithm. The flow phantom model was used in a previous study [22], where the details of manufacturing procedure were provided. The ultrasound contrast agent (DEFINITY®, Lantheus Medical Imaging, Inc.) was diluted 1000-fold with 0.9% saline and then perfused through the flow channel. The estimated average speed of the MB flow was around 8.4 mm/s by using a volume rate of 120 μL/min in the flow channel with the approximate diameter of 550 μm.

The ultrasound signal was acquired using a Verasonics Vantage 256 System (Verasonics, Kirkland, WA, USA). A 128-element high-frequency linear array transducer L35-16vX (Verasonics, Kirkland, WA, USA) was used, operating at 20MHz center frequency. Plane-wave compounding, using steering angles from −4 ° to 4 ° with a step size of 1 °, was performed to acquire signal at a post-compounding frame rate of 1,000 Hz. A total of 21 acquisitions were gathered from the CAM, with each acquisition including 1,600 frames of in-phase quadrature (IQ) data. The ULM image reconstruction was implemented as in previous studies [8,13,16]. A singular value decomposition (SVD) filter was first applied to the IQ data to filter out the MB signal by removing the low motion tissue signal and background noise [8]. A pre-acquired point spread function (PSF) of a single MB was used to calculate the 2D normalized cross correlation with each group of MB signal. An empirically chosen threshold was then used to reject small values in the correlation coefficient map and the centroids of the remaining sub-regions were used as the MB localizations. Velocity maps were generated using a fast MB pairing and tracking algorithm [46].

CTSP was compared with conventional ULM velocity reconstruction using 1.6s, 3.2s and 4.8s of acquisition data. The results were also compared with a reference velocity map generated using 33.6s of MB acquisition.

### D. In-vivo CAM study

An *in-vivo* chicken embryo CAM microvessel model was also used to validate the proposed CTSP algorithm. Fertilized chicken eggs were provided by the Poultry Research Farm at University of Illinois and placed in an incubator (Digital Sportsman Cabinet Incubator 1502, GQF manufacturing Inc.). On the fourth day, the egg shells were opened with a rotatory Dremel tool and the contents were transferred into weigh boats. Then the chicken embryos were placed into another humidified incubator (Darwin Chambers HH09-DA) until ultrasound imaging on the eighteenth day.

For the MB injection, a glass needle was made by using a PC-100 glass puller (Narishige, Setagaya City, Japan) to pull a borosilicate glass tube (B120-69-10, Sutter Instruments, Novato, CA, USA). The glass needle was then attached to Tygon R-3603 laboratory tubing. A MB solution was made with Vevo Micromarker (FUJIFILM VisualSonics) reconstituted with 1mL saline, yielding a concentration of 2 × 10^9^ MBs/mL. A total volume of 70 μL solution was injected into the embryo for ULM imaging.

The optical image of the chicken embryo was acquired as the reference ground truth image. A Nikon SMZ800 stereomicroscope (Nikon, Tokyo, Japan) with a DS-Fi3 digital microscope camera (5.9-megapixel CMOS image sensor, Nikon) was used to acquire the optical image.

The ultrasound scanning sequence was the same as what was used in the flow phantom study. In the ULM processing, the MB signal with different speed ranges and directions were separated into three groups using 3D Fourier domain filters and processed separately [16]. Velocity maps were generated using a bipartite graph-based MB pairing and tracking algorithm [8]. The final localization and velocity images were the combination of the individual reconstruction images generated from each acquisition.

To validate the CTSP, we used IQ data with data acquisition times ranging from 50ms (50 frames) to 12.8s (12,800 frames), with a step size of 50ms, to generate separate MB localization images (i.e., MB location accumulations only without MB tracking). The CTSP algorithm was applied to each 50ms data subset to recover the corrupted vessel images, which were then accumulated to generate the final CTSP recovered image. *VF_Ppercentage_* and *VF_precision_* were measured on both the MB localization images and CTSP reconstructed images with different acquisition times using both the MB localization using long acquisition (33.6s) and the optical image as ground truth. The intensity profiles of three vessels with different sizes were measured to evaluate the performance of CTSP recovery. In addition, a two-phase exponential wash-in function was used to fit a saturation curve for both datasets:

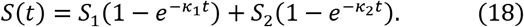

Here *S*(*t*) represents the saturation level with acquisition time, *t* is the acquisition with unit of seconds. *S*_1_ and *κ*_1_ in Eq. (18) characterize the saturation rate and the maximum level of saturation of larger vessels. *S*_2_ and *κ*_2_ in the second term characterize a much slower secondary saturation in smaller vessels with slower MB perfusion. The total maximum saturating level *S_m_* is defined as *S_m_* = *S*_1_ + *S*_2_. CTSP-recovered ULM images are expected to have significantly higher *κ*_1_ as compared with conventional localization since CTSP provides a much faster filling rate for larger vessels. CTSP is expected to have a similar *κ*_2_ as conventional localization due to physiological constraints of slow MB perfusion in smaller vessels. The 90% saturation [12–13] was used to evaluate the performance of the proposed algorithm. In addition to MB localization for vascular structure reconstruction, the CTSP was also used to recover the ULM velocity maps using 1.6s, 4.8s, and 8s of acquisition data. The results were compared with ULM velocity maps generated with a conventional MB tracking algorithm [8] using 33.6s of MB data.

### E. In-vivo mouse brain study

The mouse brain dataset used to test CTSP was obtained from a previous study [45], where the details of the animal procedure were provided. All the experiments were approved by the Institutional Animal Care and Use Committee (IACUC) at the University of Illinois Urbana-Champaign. Mice were housed in an animal care facility approved by the Association for Assessment and Accreditation of Laboratory Animal Care. A cranial window was opened on the skull of the mouse using a rotary Dremel tool for ultrasound imaging. The surgery was implemented under anesthesia using 4% isoflurane mixed with medical oxygen and 1% lidocaine applied intradermally to the scalp.

As with the *in-vivo* chicken embryo study, a Verasonics Vantage 256 System and a L35-16vX transducer were used to acquire the ultrasound data. The transducer was positioned approximately 3mm caudal to bregma and situated to make an imaging field of view that covered the entire coronal hemisphere of the mouse brain. Imaging was performed using plane-wave compounding with steering angles from −4° to 4° with a step size of 1° at a post-compounding frame rate of 1,000 Hz. A total of 81 acquisitions of IQ data were acquired, where one acquisition includes 1,600 frames (1.6s) of data. A 50 μL bolus of ultrasound contrast agent (DEFINITY®, Lantheus Medical Imaging, Inc.) was injected into the mouse tail vein for every 10 acquisitions. The ULM signal processing was similar to that used in the chicken embryo study. The microbubble pairing and tracking was performed using a fast-tracking algorithm [46].

Similarly, IQ data with acquisition times ranging from 50ms (50 frames) to 12.8s (12,800 frames), with a step size of 50ms, was used to generate localization images and for CTSP reconstruction. The final CTSP recovered image was obtained by accumulating all the individually recovered vessel images and applying a 2×2 median filter. Velocity maps using 3.2s, 6.4s, and 12.8s of acquisition data were recovered by CTSP and compared with the velocity map generated from all 81 acquisitions (129.6s).

## III. Results

### A. In-silico study

Figure 3 (a-c) shows a simulated vessel map, a randomly downsampled version of this vessel map (following a Poisson distribution) with 80.96% of the pixels removed, and the CTSP recovered image from this downsampled example. The corresponding curvelet coefficients of Fig. 3(a-c) are shown in Fig. 3(d-f), where the curvelet transform parameters (*j, l*), as described in Eq. (6), were selected as (0,0), (1,16), and (2,32). This corresponds to three layers with different scales from inner to outer in the illustration in Fig. 1(a). In Fig. 3(e), the additional high frequency coefficients caused by the discontinuities between vessel segments can be seen over all scales and orientations in the curvelet domain. After CTSP recovery, Fig. 3(f) shows significant shrinkage of the additional high frequency components, while the major coefficients that represent the vessel structure remain the same. Figure 3(c) shows the CTSP recovery in the spatial domain, where the discontinuities between vessel segments are filled in and the vessel shape is correctly recovered.

Figure 4 (a) shows the simulated microvessel maps using different amounts of acquisition time, where every vessel pixel follows the Poisson distribution with an expected value of 0.05 × *T* (where *T* represents a unit of acquisition time). Figure 4 (b) shows the CTSP recovery corresponding to the microvessel maps in Fig. 4(a). As described in Eqs. (14-15), we used an adaptive method to select the parameter updating functions *L*_1_ and *L_2_* for microvessel maps with different initial pixel densities. Six sets of parameter updating functions were first empirically selected for CTSP recovered microvessel maps using 1, 2, 4, 10, 15, and 20 units of acquisition time. Then according to Eq. (16), the regression parameters were calculated as: *L*_1_: *p*_1_ = 8.3e^-4^, *p*_2_ = –0.73, *p*_3_ = –0.009; *L*_2_: *P*_1_ = 0.42, *p*_2_ = 0.12, *p*_3_ = –0.18. Figure 5 shows the *VF_percentage_* measurement of the incomplete microvessel map and the CTSP recovery according to Eq. (17). Using 1 unit of acquisition time, CTSP can recover 86.94% of the true vessels, which is a significant improvement over the simulated accumulation microvessel map. In addition, CTSP was able to recover nearly 100% of the true vessels using 7 units of acquisition time, while the simulated accumulation microvessel map needed 100 units of acquisition time to achieve the same *VF_percentage_*, indicating a more than 14-fold increase of imaging speed. The precision measurement shows an approximate 25% of false recovered vessels on average, which was largely caused by the blurring near the vessel boundaries.

**Figure 4.**
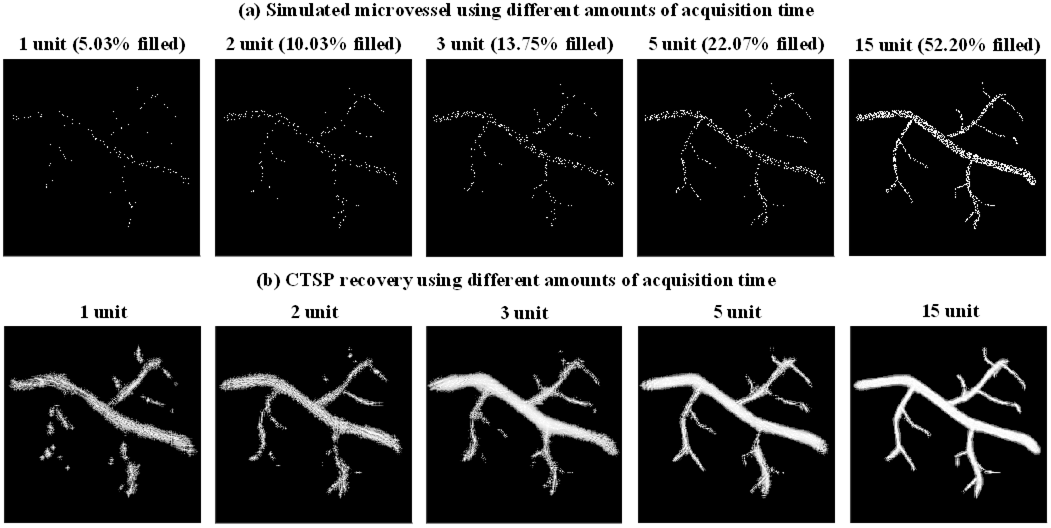
(a) Simulated microvessel image using different acquisition times (corresponding to different vessel filling percentages); (b) Corresponding CTSP recovered image from (a).

**Figure 5.**
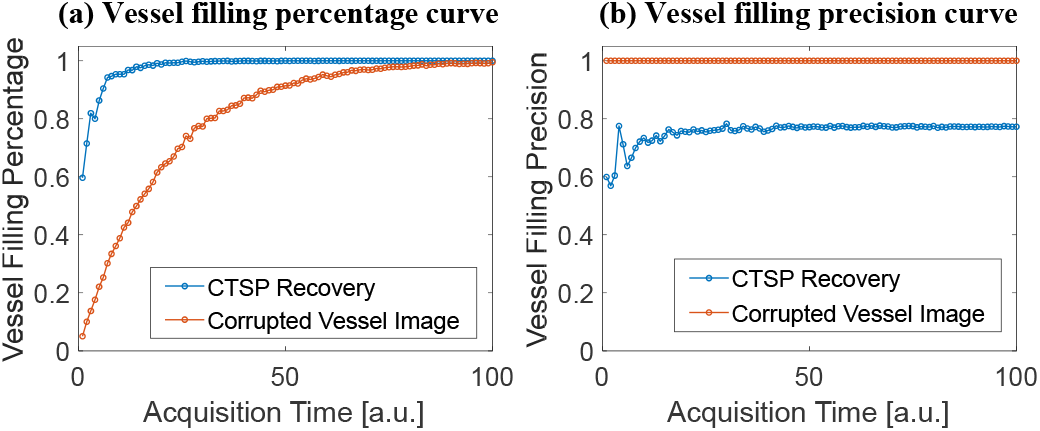
Evaluation of the CTSP performance using different metrics in the *in-silico* study. (a) The vessel filling percentage and (b) vessel filling precision of CTSP recovery and the corrupted vessel image with different acquisition times.

### B. In-vitro flow phantom study

Figure 6 shows the velocity map using MB tracking and CTSP recovery with different acquisition times. Figure 7 shows the velocity profiles of the selected vessel location in Figure 6. Three sets of data with acquisition times of 1.6s, 3.2s, and 4.8s were used to reconstruct each velocity map, with a reference image reconstructed from 33.6s of MB data. It can be seen that the velocity map reconstructed using conventional MB tracking method with short acquisition has inaccurate velocity estimation and large gaps between velocity trajectories. The velocity map recovered by CTSP shows much smoother velocity profile and more accurate flow speed estimation using less data acquisition time.

**Figure 6.**
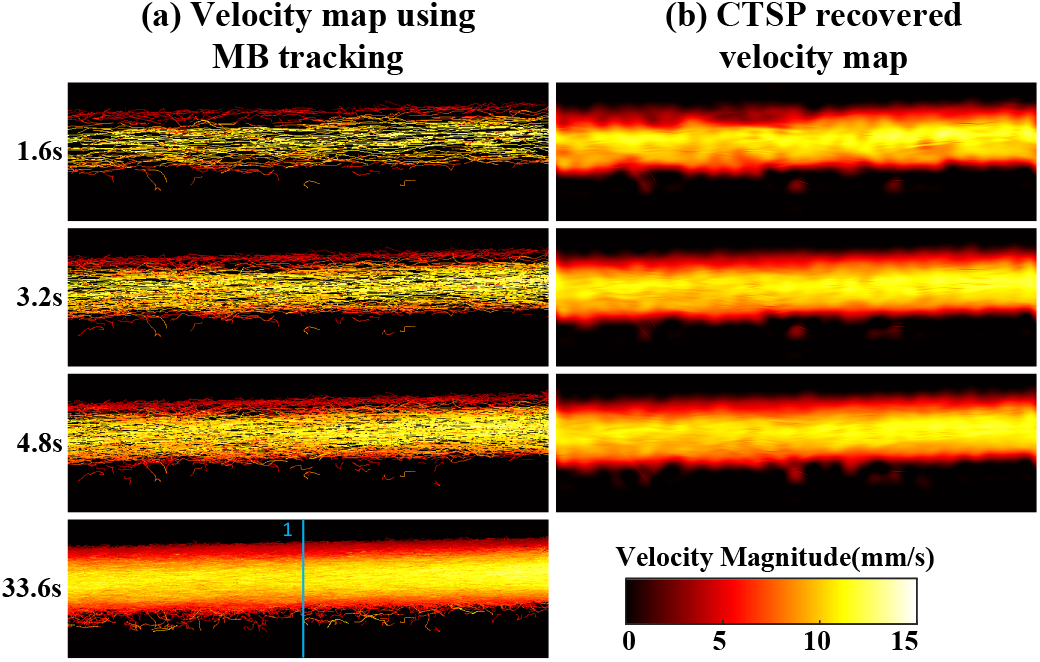
(a) Velocity maps of flow phantom using data with different acquisition times, (b) corresponding CTSP recovered velocity maps from (a). The flow direction is from the left side to the right side of the image. All the velocity maps used the identical colormap displaying velocity magnitude ranging from 0-15 mm/s. The blue line in velocity map using 33.6s acquisition marks the velocity profiles shown in Fig. 7.

**Figure 7.**
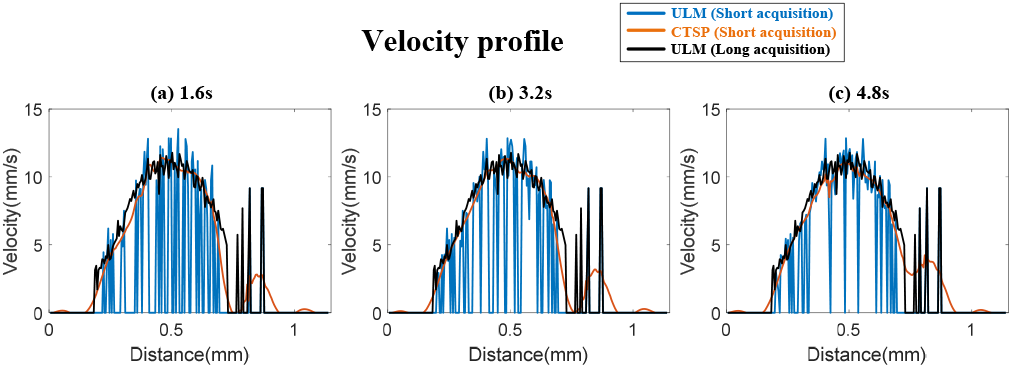
Velocity profiles of the line segment marked by the blue line in Fig. 6 (a) using 1.6s acquisition (b) using 3.2s acquisition (c) using 4.8s acquisition.

### C. In-vivo CAM study

Figure 8 shows an optical image, ULM images, and the corresponding CTSP recovered vessel images of a chicken embryo CAM. The gaps in the small vessel trajectories of the short data acquisition localization map in Fig. 8(a) could be successfully filled using CTSP recovery as seen in Fig. 8(b). The vessel structure in the CTSP recovered image was also preserved as compared to the long data acquisition time accumulation map in Fig. 8(c). Using the optical image as the reference ground truth (Figs. 8(d-e)), the CTSP recovered image using 100ms of acquisition (Fig. 8(b)) recovered 53.32% of the vessels, while the original ULM image without CTSP (Figs. 8 (a)) only revealed 5.16% of the vasculature. Even with 5 times more data (1.6s), the filling rate was still only at 32.11% (Fig. 8(c)), which is still significantly lower than CTSP with shorter data acquisition. This result demonstrates that CTSP provides a significantly faster vessel filling speed of major vessels using small number of MB events (e.g., a 100ms acquisition).

**Figure 8.**
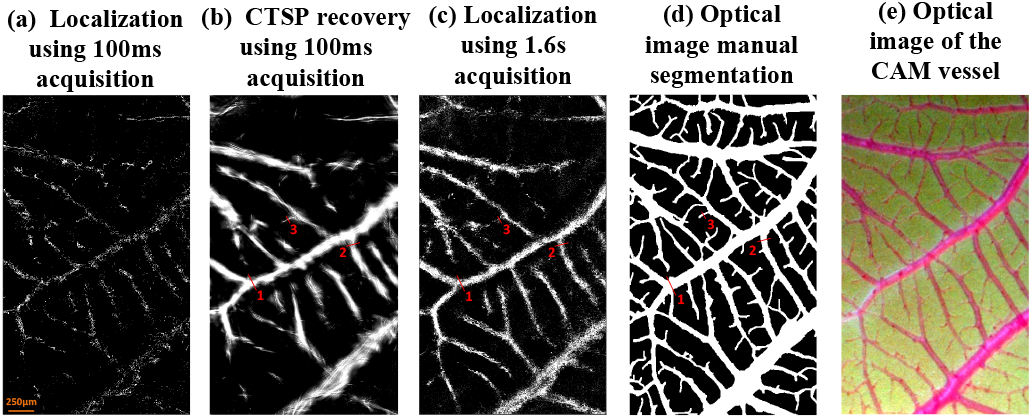
*In-vivo* optical and ULM image of a CAM vessel: localization images using (a) 100 ms (100 frames) of data acquisition; (c) 1.6s (1600 frames) of data acquisition; (b) CTSP recovered vessel image based on (a); (d) reference optical image segmentation; and (e) the original optical image. The three vessel profiles marked by red lines in (b-d) were used for evaluating the vessel width in Fig. 9.

Figure 9 shows the vessel intensity profiles of the three vessels marked by the red lines in Fig. 8. It can be seen that CTSP reconstruction using short acquisition (250ms) already had similar intensity profiles as compared with those obtained from localization using long acquisition time (33.6s). In addition, the *VF_percentage_* in Fig. 10(a) shows that using optical image as the ground truth, the maximum saturation level *S_m_* as described in Eq. (18) is 67.20% for MB localization and 85.34% for the CTSP recovery. The 90% saturation time of the MB localization is 2.4s, while CTSP only needs 0.2s to achieve the same level of saturation, indicating a 12-fold faster imaging speed. In addition, CTSP demonstrates the ability to recover the major vessels (42.2% of the true vessels) using a very short acquisition time (50ms), while MB localization recovered 2.7% using the same 50ms acquisition time. Similar level of improvement can be also seen in Fig. 10(b) using long acquisition localization as the ground truth. In Fig. 10(c), the precision measurement using optical image as ground truth shows about 50% false positive rate in both CTSP recovery and localization images. This level of false positive rate was largely caused by the misalignment between the optical and ultrasound image. As shown in Fig. 10(d), the false recovery rate dropped to 20-30% when CTSP was compared with MB localization using long data acquisition (33.6s), which was similar to the simulation results shown in Fig. 5(b). Figure 11 shows ULM images of the same local region in Fig. 8 of the CAM vessel using different data acquisition times along with the corresponding CTSP recovered images. Six sets of parameter updating functions were first selected for CTSP recovery using 50, 100, 150, 500, 1000, and 1500ms of acquisition time. Then the regression parameters were calculated for the adaptive parameter updating function as: *L*_1_: *p*_1_ = 1.56e^-4^, *p*_2_ = – 1.26, *p*_3_ = –0.0023; *L*_2_: *P*_1_ = 0.35, *p*_2_ = 0.60, *p*_3_ = 0.01.

**Figure 9.**
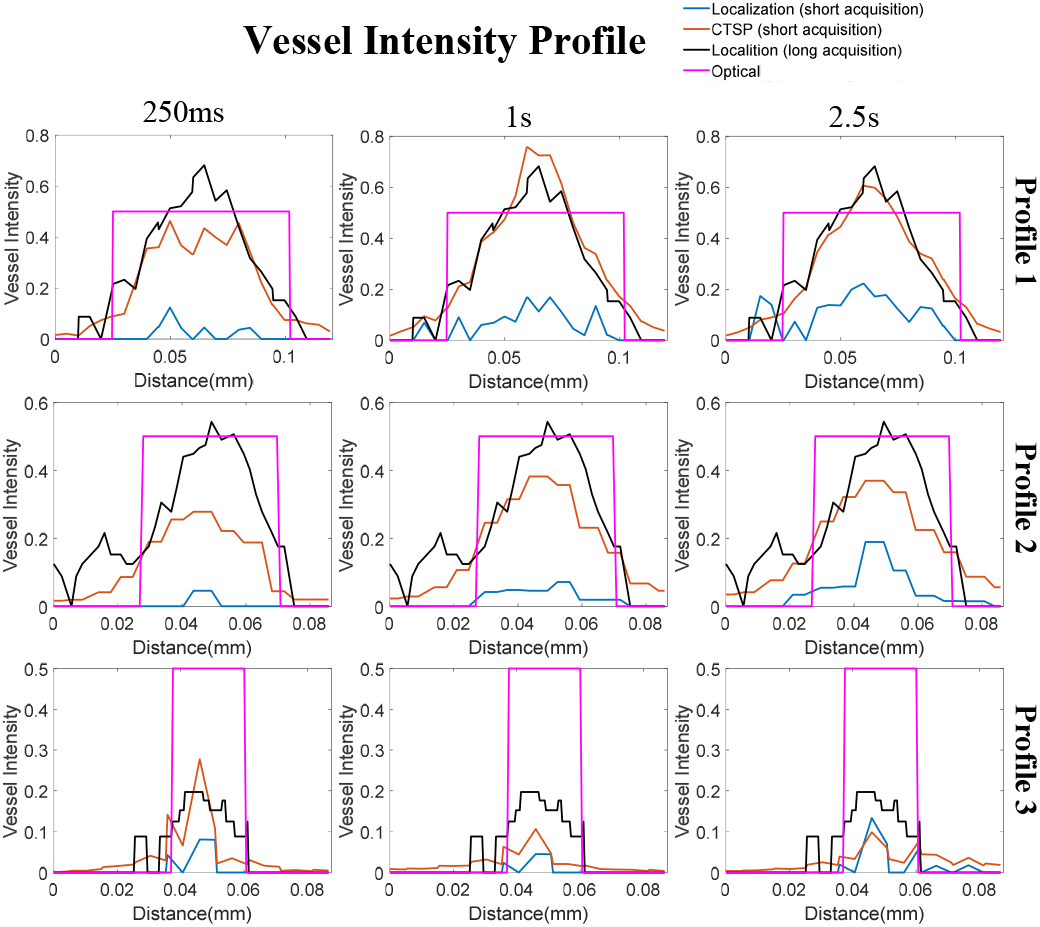
Vessel intensity profiles of the three vessels marked in Fig. 8 (b-d). The three columns represent the different acquisition times used for reconstruction. The optical segmentation image and the long accumulation (33.6s) ULM image were used as ground truth. The vessel profile based on the optical segmentation image (purple) was set at 0.5 intensity value for better visual comparison.

**Figure 10.**
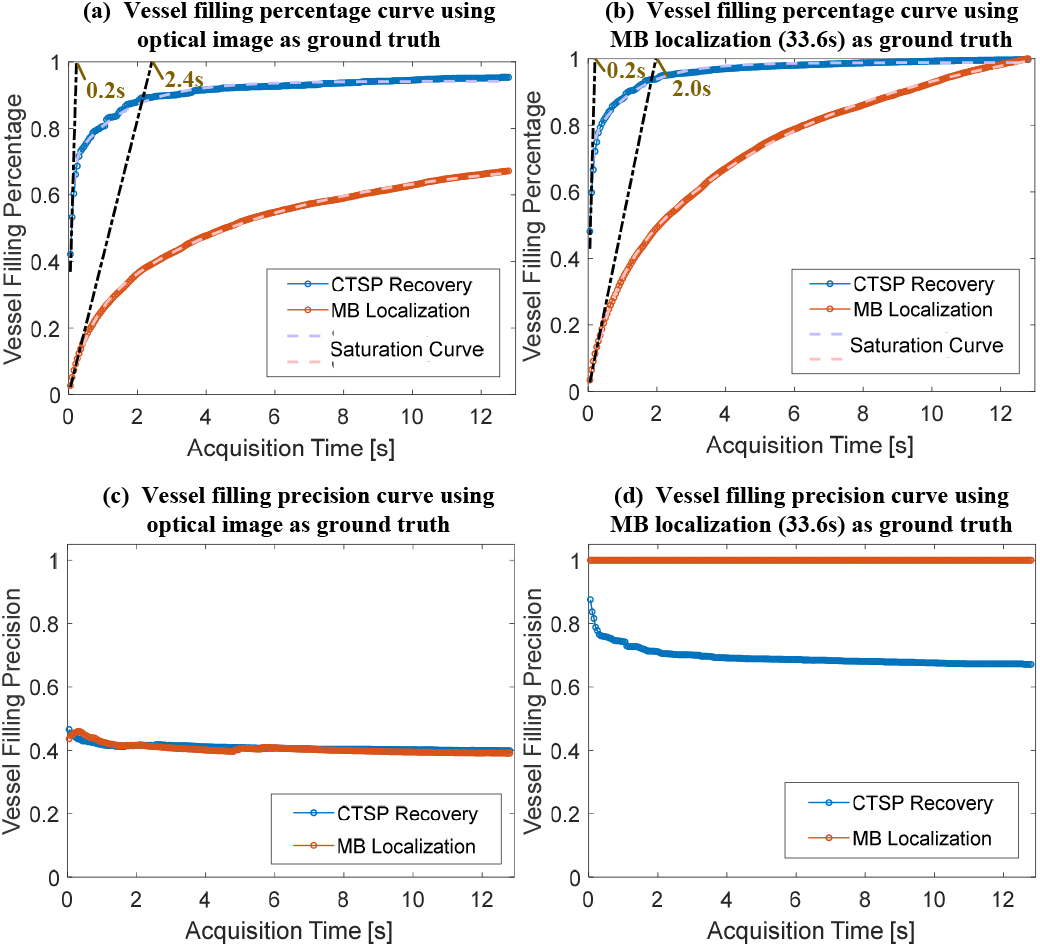
Evaluation of the CTSP performance using different metrics in the *in-vivo* CAM study. The vessel filling percentage with different data acquisition times (a) using optical image as ground truth (b) using MB localization (33.6s) as ground truth. The vessel filling precision curve with different data acquisition times (c) using optical image as ground truth (d) using MB localization (33.6s) as ground truth.

**Figure 11.**
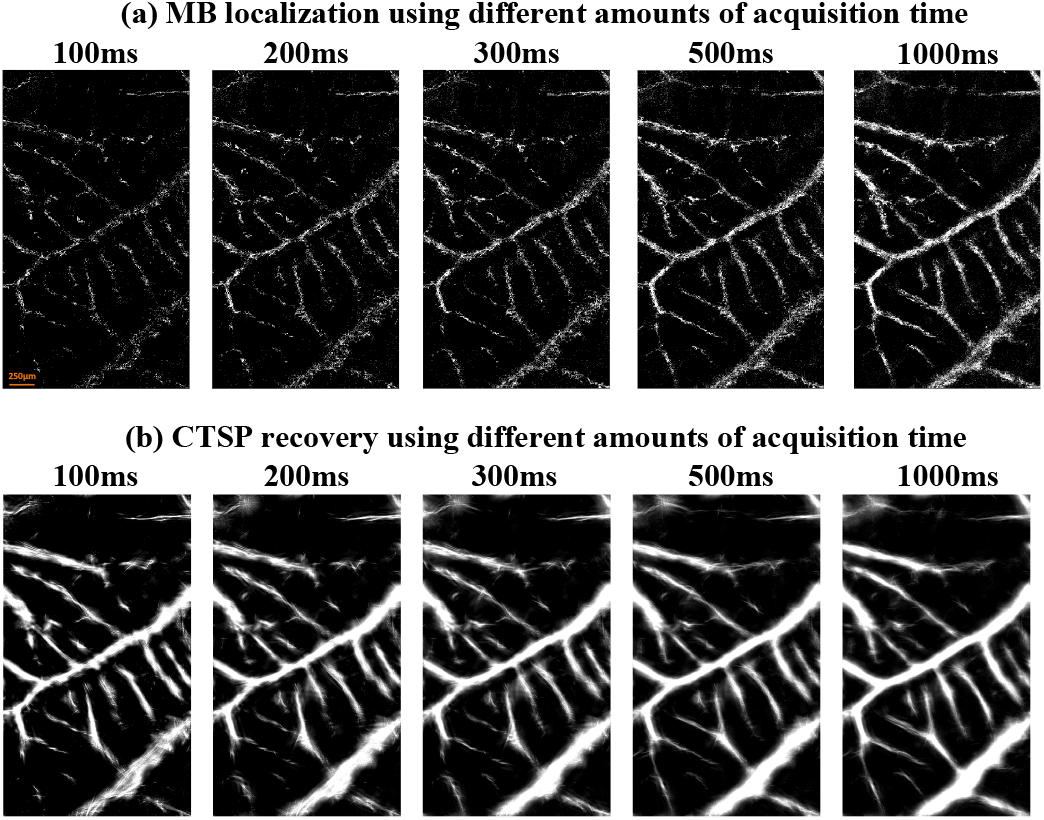
(a) *In-vivo* ULM image of a CAM vessel using data of different acquisition times (frame rate = 1000 Hz); (b) Corresponding CTSP recovered image from (a).

Figure 12 shows velocity maps of CAM vasculature reconstructed via conventional MB tracking and the corresponding CTSP recovered results. Three data subsets with acquisition times of 1.6s, 4.8s, and 8s, were used to reconstruct velocity maps. A reference image was reconstructed with 33.6s of acquisition time. As shown in Fig. 12(a), with 1,600 frames (1.6s) of data, the velocity map generated by MB tracking is very noisy, which is likely due to unreliable flow speed estimation from using limited number of MB localization samples. As the amount of data increases, the velocity maps become smoother with less fluctuations within local regions of the vessel. The velocity measurement fluctuations caused by insufficient MB data also lead to additional high frequency components in the curvelet domain, as shown in Fig. 14(a,d). Therefore, by suppressing the excessive high frequency components in the curvelet domain, as shown in Fig. 14(b,e), CTSP was able to remove the noise in vessel flow velocity estimation and restore high quality flow velocity maps. It can be seen from Fig. 12(b) that after CTSP recovery, the velocity map becomes smoother and is closer to the reference image shown in Fig. 12(a). In addition, as shown in Fig. 13, CTSP was able to reconstruct the vessel cross-section velocity profiles to nearly identical with that from the 33.6s reference acquisition, despite only using only 4.8s of acquisition time.

**Figure 12.**
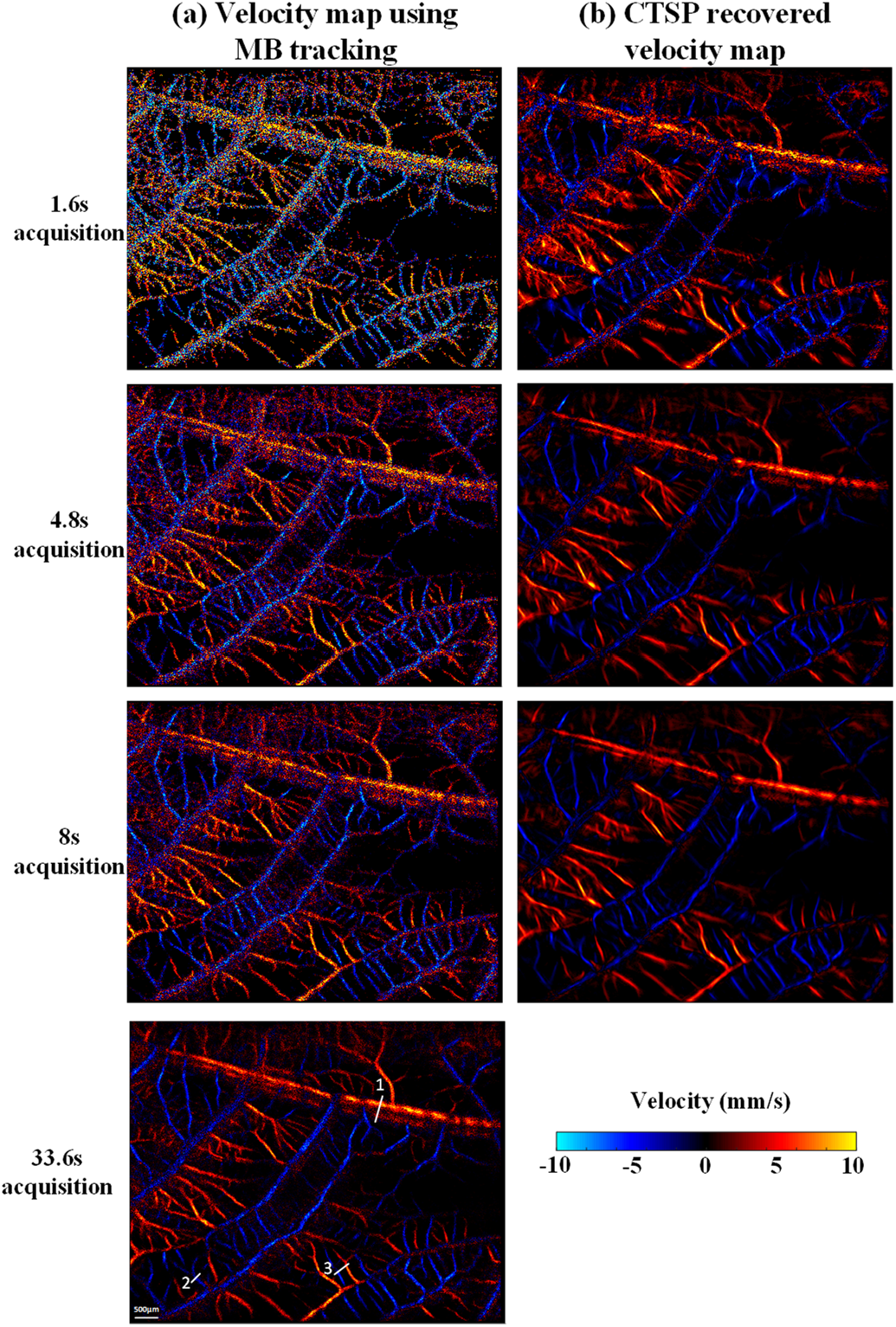
(a) Velocity maps of CAM vasculature using data with different acquisition times, (b) corresponding CTSP recovered velocity maps from (a). All the velocity maps use the identical colormap with the velocity range from −10 to 10 mm/s. The three vessel profiles marked by white lines in are used for evaluating the velocity recovery in Fig. 13.

**Figure 13.**
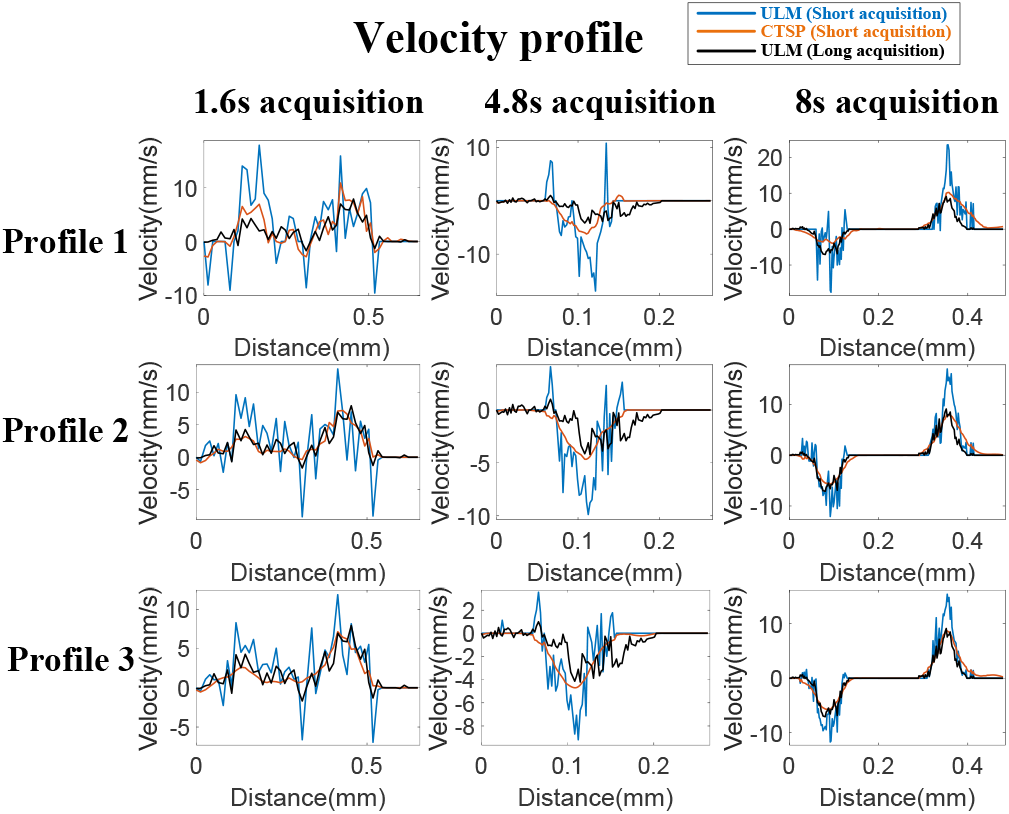
Velocity profiles of the line segments marked by the white line in Fig. 12 using different acquisition time.

**Figure 14.**
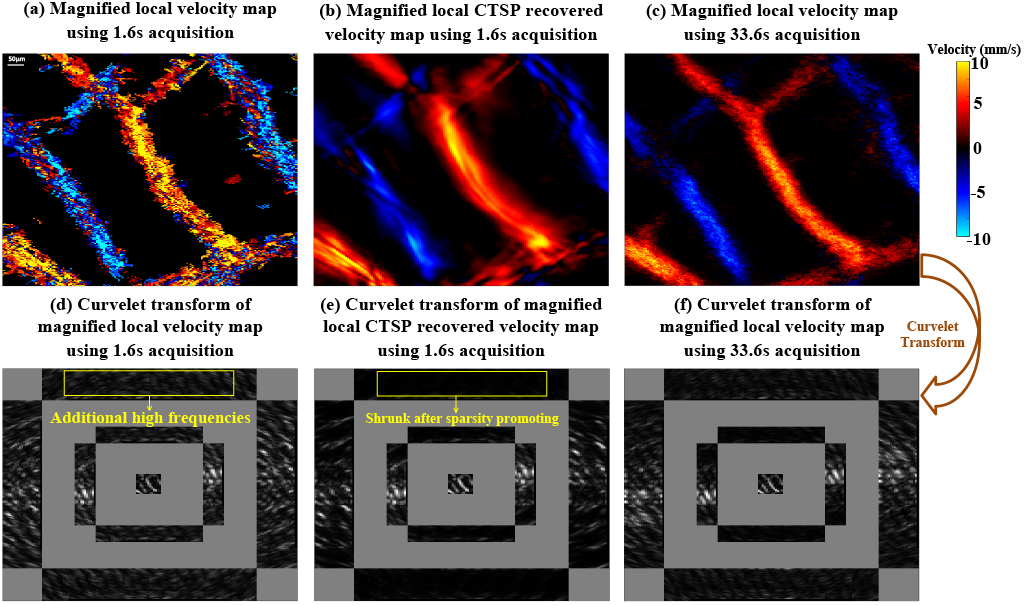
Velocity maps of a local region of CAM vasculature from Fig. 12 (a) MB-tracking using 1.6s acquisition time; (b) CTSP recovery using 1.6s acquisition time; (c) MB-tracking using 33.6s acquisition time, (d-f) corresponding curvelet transforms of the images in (a-c). All the velocity maps use the identical colormap displayed in (c) with the velocity range from −10 to 10 mm/s.

### D. In-vivo mouse brain study

Figure 15 shows the ULM images of a mouse brain using different data acquisition times and the corresponding CTSP recovered images. Similar to the chicken embryo CAM study, ten sets of parameter updating functions were empirically chosen for CTSP recovery using 100, 200, 300, 700, 1000, 1500, 2500, 5000, 7500, and 12500ms of acquisition times. The regression parameters were calculated for the adaptive parameter updating function as: *L*_1_: *p*_1_ = 5.18e^-4^, *p*_2_ = −0.97, *p*_3_ = –5.76c^-4^; *L*_2_: *P*_1_ = 0.23, *p*_2_ = 0.44, *p*_3_ = 0.03. Fig. 15(b) demonstrates that CTSP can connect the large vessel branches from the sparse MB locations in Fig. 15(a) using only 500ms of acquisition data. In Fig. 15(b), it is shown that CTSP can reconstruct most of the vasculature using just 1-2s of acquisition data, with MB localization image using 129.6s as a reference.

**Figure 15.**
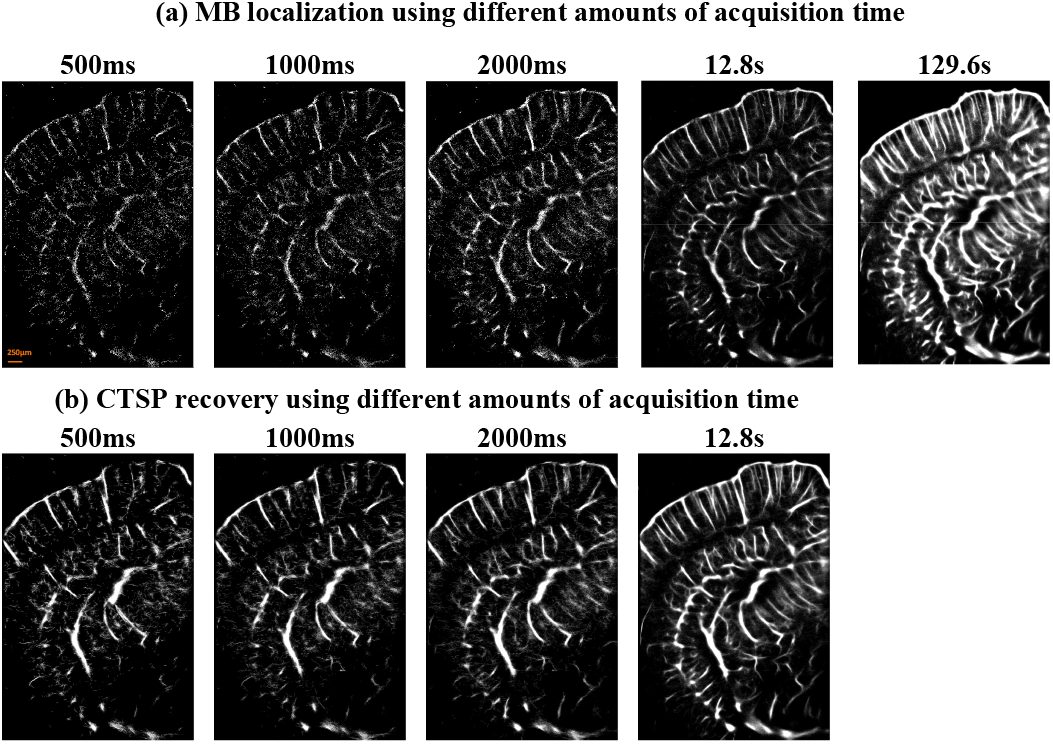
(a) *In-vivo* ULM image of a mouse brain vessel using data of different acquisition time (frame rate = 1000 Hz), (b) corresponding CTSP recovered image from (a).

Figure 16 shows the velocity maps of the mouse brain from using conventional MB-tracking and the corresponding CTSP recovered velocity maps. Three sets of data with acquisition times of 3.2s, 6.4s, and 9.6s were used to reconstruct each velocity map, with a reference image reconstructed with a total of 129.6s of data. Similar to the velocity map of the CAM, the velocity map generated by MB tracking using an inadequate amount of data (e.g., 3.2s −9.6s acquisitions) lacks precision on the velocity estimations. CTSP can recover the velocity value by removing additional high frequency components and promoting the sparsity in the curvelet domain. It can be seen from Fig. 16(b) that CTSP recovery can recover the velocity maps in Fig. 16(a) and make the velocity values closer to the reference velocity image generated using 129.6s of acquisition, as shown in Fig.16(a). In addition, the velocity profiles in Fig. 17 show that CTSP can correct the inaccurate velocity values caused by a short data acquisition time. These results show that CTSP is able to reconstruct high-fidelity ULM velocity maps with much less MB data than conventional tracking algorithms, leading to significantly reduced imaging time and subsequently higher temporal resolution for ULM.

**Figure 16.**
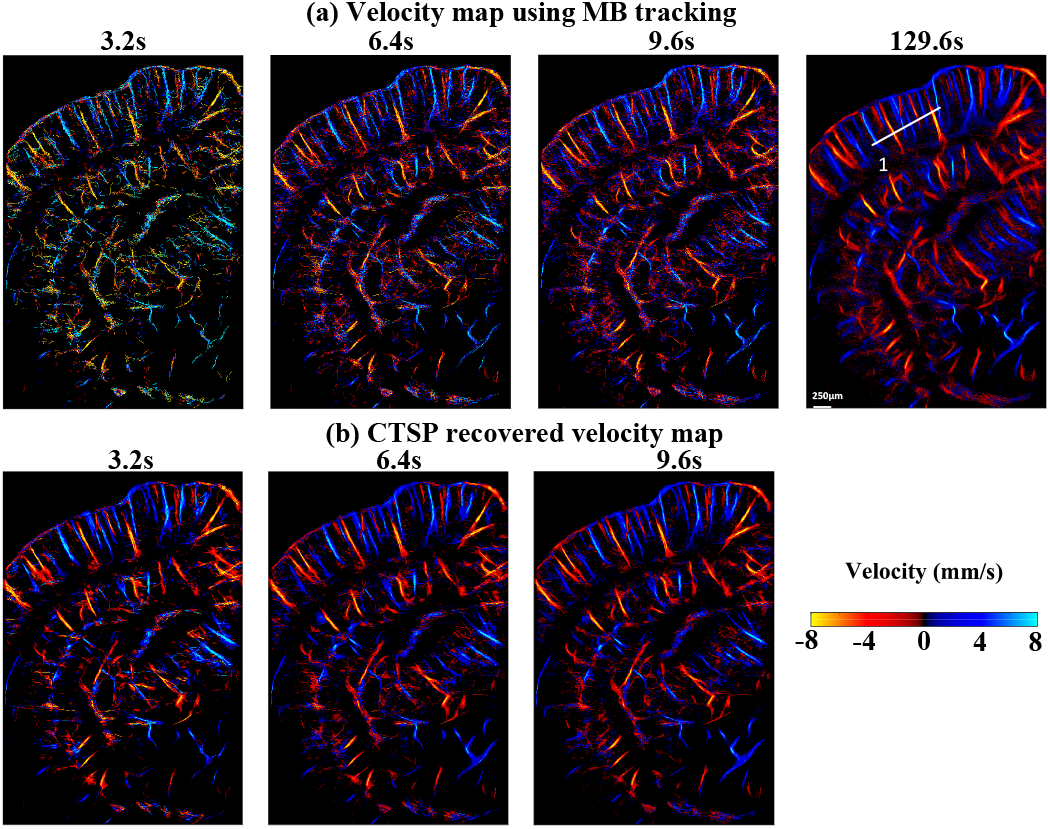
(a) Velocity maps of a mouse brain vessel using data of different acquisition time, (b) corresponding CTSP recovered velocity maps from (a). All the velocity maps use the identical colormap displayed in (c) with the velocity range from −8 to 8 mm/s. one vessel profile marked by white lines in are used for evaluating the velocity recovery in Fig. 17.

**Figure 17.**
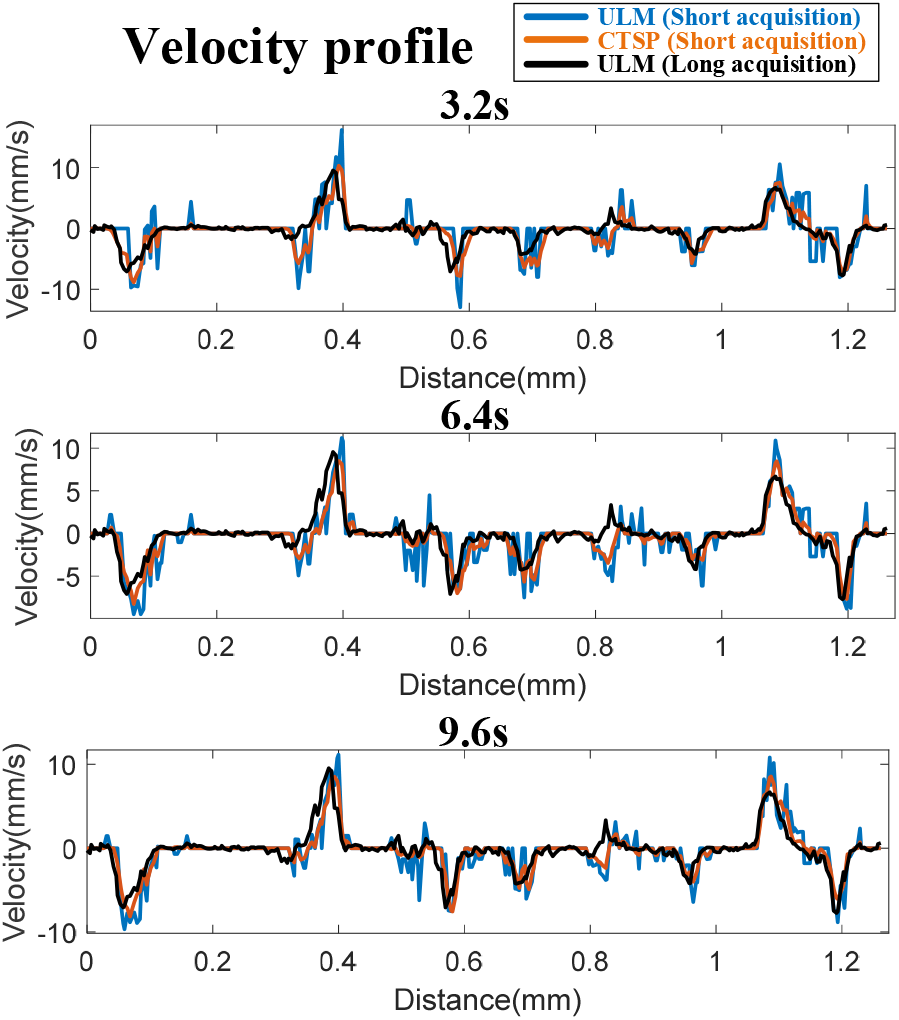
Velocity profile of the line segment marked by the white line in Fig. 16 using different acquisition time.

## IV. Discussion

This paper presented a curvelet transform-based sparsity promoting (CTSP) algorithm for fast ultrasound localization microscopy (ULM) imaging. The proposed method uses a compressive sampling model to describe the problem of microvessel reconstruction from sparse MB localizations. In addition, the proposed CTSP method was based on several characteristics of ULM imaging: first, ULM imaging reconstructs microvessel structure by accumulating MB locations in the blood flow under the assumption that MBs do not extravasate from the blood pool and can mimic the rheology of red blood cells; two features that are well established in the literatures [3–8]. In addition, the relatively low MB concentration in the blood implies that each detected MB location at a fixed time point can be considered as a random spatial sampling of the true vessel location(s). Second, the microvessel structure can be considered sparse in both the curvelet domain and the spatial domain, which is substantiated by the low reported vascular volume fractions in tissues [23]. This sparsity guarantees the possibility of recovering the microvessel structure from an inadequate amount of MB locations. Under these axioms, CTSP is theoretically able to recover the microvessel structure from sparse spatial signal including MB locations and/or MB tracking trajectories. Therefore, the proposed method can be applied to any MB localization or tracking techniques that are using an inadequate amount of MB signal. In our study, it was demonstrated that CTSP can achieve a faster and more accurate reconstruction of the microvessel structure than conventional ULM techniques in both simulation and *in-vivo* experimental data.

The presented results showed that CTSP has several distinct advantages over existing ULM strategies. First, CTSP recovers the major branches of the microvasculature using very little MB signal. This was demonstrated in the chicken embryo CAM vasculature, where CTSP could recover the principal vessels from very sparse MB distributions accumulated in just 50 frames (50ms of acquisition time), which corresponds to a 48.1% vessel filling rate (Fig. 10). Using 200ms or 300ms acquisition time, the smallest vessels that can be resolved from CTSP recovered images have a width of around 20-50 μm (Fig. 11). In the mouse brain study shown in Fig. 15, CTSP could reconstruct most of the vessels in the whole brain cross-section using just 500 to 1000 frames of MB data (0.5s to 1s of acquisition time). The vasculature in the mouse brain is more complex than that in the chicken embryo CAM, therefore a longer acquisition time was necessary to review the entire vasculature. One strategy to leverage the high imaging speed of CTSP recovery is to use it as a scouting technique to confirm the targeted tissue vasculature before executing a long data acquisition sequence for ULM. For example, short test acquisitions of MB data can be quickly reconstructed using CTSP to reveal the tissue microvasculature, which could aid in confirming the placement of the ultrasound transducer to best capture the region of interest. CTSP is also more efficient at utilizing the spatially sparse MB localizations present in small vessels that may otherwise need a large amount of data acquisition time to completely scout using conventional localization techniques. This feature is most evident in the saturation curves presented in Fig. 10, where CTSP increased the maximum saturation level of MB accumulation data relative to the optical imaging reference on CAM microvasculature - the saturation curve of CTSP recovery converged to 90% of the microvessels present in the optical image while the saturation curve of localization reconstructed 67% of the vessels in the reference. In the simulation study (Fig. 5), CTSP very rapidly reconstructed the majority of the vasculature in the first few time steps and then proceeded at a much slower rate in comparison to the more gradual perfusion evident in conventional ULM accumulation. We posit that the gain which CTSP exhibits in the saturation curves is due, in part, to a compensation for perfusion statistics: CTSP can efficiently recover sparse MB events in well perfused vasculature, which accelerates the early phase of saturation, but the late phase is still dictated by the physiological constraints of rarely perfused microvessels [12].

Secondly, CTSP has also shown the ability to recover velocity map information using MB data with a short acquisition time due to the signal denoising effect after curvelet sparsity promotion. Typically, an accurate estimation of blood flow velocity requires a large amount of MB signal [7] to fully characterize the flow velocity profiles. As shown in Fig. 14, the velocity map estimated using an inadequate amount of MB signal has noisy velocity measurements and cross-section profiles. In the curvelet domain, these velocity fluctuations lead to additional high frequency components that are not present in the curvelet transform of the reference velocity map. CTSP can effectively remove these additional frequency components by promoting sparsity in the curvelet domain, which generates velocity maps that have similar qualities as the reference image. The improvement in ULM velocity map reconstructions based on CTSP was validated on both chicken embryo CAM (Fig. 12) and mouse brain studies (Fig. 16).

Finally, it is worth noting that the computational cost of CTSP is primarily associated with the method used to solve the optimization problem in Eq. (9). We followed previous studies [26,29] and implemented the block-coordinate-relaxation (BCR) method to solve Eq. (9). The BCR method normally requires a large number of iterations to converge, and thus a regularization parameter updating strategy is also needed to improve the converging speed. Herrmann et al. [26] initialized the regularization parameter as the maximum value in the sparsifying domain and then used a decreasing function for updating. Elad et al. [29] designed the regularization parameter updating function to linearly decrease with the iteration index as described in Eq. (14). Both studies required dozens to hundreds of iterations to generate their results, and these updating functions have the significant drawback of a lack of flexibility for different initiation signal. In our study, the initial ULM images using data from different acquisition times had different vessel densities in both the spatial and the curvelet domain. Therefore, an adaptive regularization parameter updating strategy, as designed in Eqs. (14-15), was introduced to achieve the optimal converging speed and reconstruction accuracy for various initiations. In both the simulation study and the *in-vivo* study, the CTSP algorithm using the proposed adaptive regularization parameter updating strategy took 10-20 iterations in total to generate the reconstruction results. In addition, the scales of the curvelet transform also needs to be carefully chosen to balance the computing cost and the performance of CTSP. In our study, we chose the number of scales as *N_scale_* = *ceil*(log_2_(min(*N*_1_, *N*_2_) – 3)) and the number of orientations as *N_orientation_* = 16 * 2^*ceil*(*N_scale_*/2)^, where *N*_1_ and *N*_2_ represent the size of the spatial image, *ceil*(*) represents the operator that takes the nearest integer greater than or equal to the input number. The number of translation k was decided by the correlation between the specific curvelet and the 2D image. These curvelet transform parameters were chosen based on the fast discrete curvelet transform proposed by Candès *et al* [42] and proved to be appropriate for the CTSP algorithm proposed in this study. Using a computing platform with a single Intel CPU (i9-10900K, 10 cores, 5.2GHz frequency), reconstructing a 1000 × 800 image with CTSP took 10-20s. Further improvement of computational speed could be expected from parallel implementation using GPUs.

One limitation in the CAM study is the comparison between optical image and ultrasound image. Because the surface of the CAM was not entirely flat, some vessels shown in the optical image of the CAM (e.g., the vessels close to the transducer surface) were out of the ultrasound imaging plane. In addition, the optical image was not acquired simultaneously with the ultrasound scanning, which leaded to the misalignment between the ultrasound image and optical image. In our study, images from these two modalities were registered using normalized cross correlation before comparison. However, it is still difficult to achieve a pixel-to-pixel alignment between optical and ultrasound images. The misalignment and missing vessels will lead to a degraded performance in the precision measurement using optical image as ground truth.

An important feature of CTSP is that the recovery of microvessel structure is only based on current MB localizations. Although CTSP can recover localization images using data from short acquisition times, it cannot generate new MB signal and locations. As can be seen in Figs. 8(b) and (d), CTSP recovery could connect most of vessel branches using 100 frames of data but was not able to recover the vessel branches near the top edge of the image. The reason for this limitation is that the compressive sampling theory applied by CTSP requires that the spatial image have a lower limit of sampling rate. In our study, when the spatial sampling of microvessel structure (MB locations) is too sparse to generate enough curvelet coefficients that contain the spatial structure information, the true vessels are no longer recoverable by CTSP, and the recovery of these vessels is dictated by rare perfusion events. In addition, under the circumstances of long MB acquisition in dense microvasculature, the assumption of sparsity is violated, which results in suboptimal CTSP recovery. Generally, CTSP has the best performance for MB signal within a specific range of acquisition time. The MB acquisition time can be empirically chosen so that the input MB density is appropriate for CTSP. In our CAM and mouse brain studies, CTSP was proved to be most effective for short MB signal acquisitions from 50ms to 100ms, which equals to the MB concentration of 600 to 3300 *counts/mm*^2^. For longer data acquisitions, we applied CTSP to short time segments (e.g., 50ms) of data ensembles and accumulated all the individually recovered CTSP images to generate the final image. Moreover, the performance of CTSP is also dependent on the fidelity of MB localization signal. For example, noise contaminated and false MB localizations can lead to compromised CTSP recovery performance. Prelocalization filters such as the non-local means filtering [8] is effective with improving the CTSP recovered image quality. In this study, we used the microbubble separation technique [16], which also improves microbubble signal quality. However, in addition to noise, false MB localization can still happen because of tissue motion, overlap MB events, and distorted PSFs. The false localizations will inevitably generate artifacts in the CTSP recovered images. Therefore, we elected to use a small median filter (2 × 2) in this study to alleviate the issue of noise and false MB localizations to facilitate robust CTSP. Based on the results the median filter was effective with the noise and false MB location removal while introducing negligible blurring to the final super-resolved ULM images.

Another drawback of CTSP is that the recovered microvessel image using short acquisition does not exactly match the images reconstructed using conventional ULM with long data acquisition. First, the shrinkage of the curvelet coefficients leads to overfilling (e.g., blurring, false positives, etc.) in the image domain, leading to false positive recoveries. In our study, CTSP consistently showed 20-30% false positive rates in regions close to the vessels. However, with the constraints from spatial sparsity and total variation regularization, false positive rates were low for non-vessel regions (e.g., the dark background between visible vessels), which is an important feature of the CTSP algorithm and essential in practice for not creating artificial vessels.

Secondly, CTSP recovery could shrink some of the curvelet coefficients that contain the information of the small vessels, which could make the small vessels undiscernible. The blurring can be controlled by choosing the optimized Lagrange multipliers *λ*_1_ and *λ*_2_ in Eq. (9) that balance the sparsity promoting in the *ℓ*_1_ norm term and the data fidelity in the *ℓ*_2_ error term. However, the tradeoff between vessel recovery and blurring still exist in the CTSP recovery and needs to be carefully balanced based on different applications. In our study, we used the adaptive parameter update function in Eq. (15) to select the best *λ*_1_ and *λ*_2_ based on visual judgment of the recovered image quality. In addition, the reconstruction parameters are always adjustable based on the requirements for the specific applications.

The above spatial feature changes brought by CTSP will commonly exist in the compressive sampling reconstruction algorithm. An appropriated regularization term (e.g., total variation) can eliminate this effect by stabilizing the uncertainty with respect to residual components in the curvelet reconstructed image [47–48]. In addition, it is essential to choose the sparsifying transform that can best represent the spatial feature of the original image. As shown in Figs. 1-2, the curvelet bases have similar shape as vessels in the spatial domain and their orientation property guarantees the sparsity in curvelet domain. Therefore, even though curvelet transform is not orthogonal transform, which means it can cause uncertainty during reconstruction [49], it still showed to be highly effective in recovering corrupted vessel and velocity map. Although the other more commonly used transforms (e.g., wavelet and 2D Fourier transform) could recover the missing vessels to a certain extent, they do not offer as good performance as CTSP. One possible reason for the suboptimal performance of wavelet and 2D Fourier transform is that neither was designed to preserve vascular morphologies, which is an important feature of the curvelet transform. Other oriented frames such as shearlets [50] may have similar properties as curvelets and also be applicable to the sparsity promoting recovery for ULM, which will be investigated in a future study.

## V. Conclusion

This paper presents a curvelet transform-based sparsity promoting (CTSP) algorithm aimed at improving the image speed of the ultrasound localization microscopy (ULM) by recovering missing MB localization signal from short data acquisitions. The proposed CTSP method consists of a compressive sampling-based ULM model and an iterative method that can effectively solve the *ℓ*_1_ sparsity optimization problem. The *in-silico* simulations, *in-vivo* chicken embryo CAM, and *in-vivo* mouse brain studies have validated that CTSP could effectively recover the missing vessel signal and markedly improve the quality of the ULM velocity maps using very short MB signal accumulations. This technique provides a robust solution for practical implementations of ULM where fast reconstructions of vascular images with short data acquisitions is essential.

